# A Machine Learning Method to Characterize Conformational Changes of Amino Acids in Proteins

**DOI:** 10.1101/2023.04.16.536913

**Authors:** Parisa Mollaei, Amir Barati Farimani

## Abstract

Amino acid dynamics are significant in determining the overall function, structure, stability, and activity of proteins. However, atomic-level descriptions of the structural features of proteins are limited by the current resolutions of experimental and computational techniques. In this study, we developed a machine learning (ML) framework for characterizing the individual aminoacids dynamic in a protein and compute its contribution to the overall function of proteins. This framewor identifies specific types of angular features in amino acids, such as bimodal-switch residues. It can assist in the analysis of various protein characteristics and provide valuable insights into the dynamic behavior of individual amino acids within a protein structure. We found that there is a strong correlation between a specific type of bimodal-switch residues and the global features in proteins. This knowledge can help us to identify key residues that are strongly correlated to the overall function of the protein.

## Introduction

Proteins are dynamic molecules, and the motions of the atoms and amino acids within each protein describes its function. These motions can range from small-scale vibrations and rotations of individual amino acids to large-scale conformational changes of entire protein domains. The dynamics of amino acids in proteins are significant because they play a crucial role in determining the overall structure, stability, activity, and function of the proteins.^1–14^ Residues can interact with one another through various types of chemical bonds, such as hydrogen and disulfide bonds, which contribute to the overall function and activity of protein.^9,10^ Many proteins undergo conformational changes in response to specific signals or stimuli, allowing them to carry out their biological functions.^15,16^ Furthermore, some proteins can be regulated by other molecules that bind at a site separate from the active site, which can cause conformational changes that alter the protein’s activity.^17–23^ However, investigating the dynamics of proteins at atomistic level poses challenges both experimentally and computationally.^4,6,24–28^ Techniques such as X-ray crystallography, NMR spectroscopy can be used to characterize the amino acids structure.^13,29,30^ However, current spatial-temporal resolutions prohibit the accurate description of biophysical features of individual amino acids.^14,31–35^ The rapid motion of amino acids occurs at frequencies in the gigahertz (GHz) range, which cannot be precisely resolved by current instrumentation. On the other hand, molecular dynamics (MD) simulations can be used to study protein conformational changes at atomic resolution with femtosecond (fs) precision.^36–39^ However, achieving statistically significant results requires a substantial number of sampling and simulations. Even if the simulation data is available, analyzing a vast number of trajectories and extracting biophysical mechanisms can still be a challenging task. To this end, knowledge of statistics and machine learning is required to deal with large amounts of data for capturing residue-level conformational changes in proteins.^40–44^ The advantage of statistics with MD simulation is utilizing numerous short trajectories of protein dynamics at different states and extracting biophysical knowledge. However, the challenge arising in this scenario is the size of the data involved, as measuring all the angles in individual amino acids across the entire protein in thousands of trajectories can provide a large amount of data. To overcome these challenges, in this study, we developed a framework that incorporates modern processing of trajectories and uses machine learning to characterize amino acid motion and their biophysical behavior. ML methods can automatically uncover hidden patterns and relationships in data, which can be used to make predictions or to cluster similar data points.^41,45,46^ This model enables us to uncover the angular characteristics of each amino acid within proteins belonging to different categories. As long as a residue undergoes no significant conformational changes, its angles tend to remain stable, resulting in a uni-modal density of states. On the other hand, during the protein’s dynamics, certain amino acids experience conformational changes, causing some angles within the residue to oscillate between two or more states. This leads to the formation of a multi-modal density of states. In this study, we have investigated a specific type of angular feature in amino acids that switch between only two states. This bimodal-switch residue undergoes a sharp jump between the two angles without passing through any intermediate states during the protein dynamics. The identification of such switch residues can assist in the analysis of various protein characteristics (such as signal transduction pathways, folding processes, conformational changes, etc) which provides valuable insights into the dynamic behavior of individual amino acids within the protein structure. By tracking such residue movements, we can identify which regions of the protein are most important for its function and how changes in these regions can lead to alterations in the protein’s overall behavior. Finally, we verified our method by applying it to Fs peptide and *β*_2_*AR* receptor trajectories and characterized the bimodal-switch residue behavior. In addition, we found that there is a strong correlation between the bimodal-switch residues and the global structural features of the protein.

## Methods

### Data preprocessing

Fig.1 describes the framework that we developed to characterize the switching behavior of residues in proteins. The framework is then demonstrated through two examples to validate the performance and effectiveness of our model. To identify the atomic-level conformation of proteins, the Protein Data Bank (PDB) is a necessary resource as it provides a database of 3D structural data for proteins.^47–49^ To begin, the molecular dynamics (MD) simulation of a protein, which is a valuable tool for obtaining information about the 3D conformational dynamics of a protein,^36,50,51^ is required in order to characterize dynamics of the switch residues. To examine the angle characteristics within a protein, we identified specific atoms located at the edges and nodes of each amino acid in order to determine significant angular features in it (see Supporting Information). This approach also aids in decreasing a large number of angles within one single residue. In that way, we could reduce 2024 angles in ARG residue to 120 important angles. By selecting these atoms within each amino acid, for example, the *β*_2_*AR* receptor (PDB 3P0G)^52^ can yield 12138 angles generated by 271 amino acids. To achieve a density of states, we measured each of these angles in every frame throughout the entire trajectory. Therefore, as shown in pre-processing section in Fig.1, the final stage involves density of states for each individual angle state. Proteins can exhibit different angle distributions that fall mainly into three categories: uni-modal, bi-modal, and multi-modal, with numerous variations within each category (some examples are shown in Fig.1 and Fig.2). The decision to utilize machine learning (ML) models for the classification of bimodal distributions was made due to the vast size of the data and the wide range of angle distributions presented.

**Figure 1:**
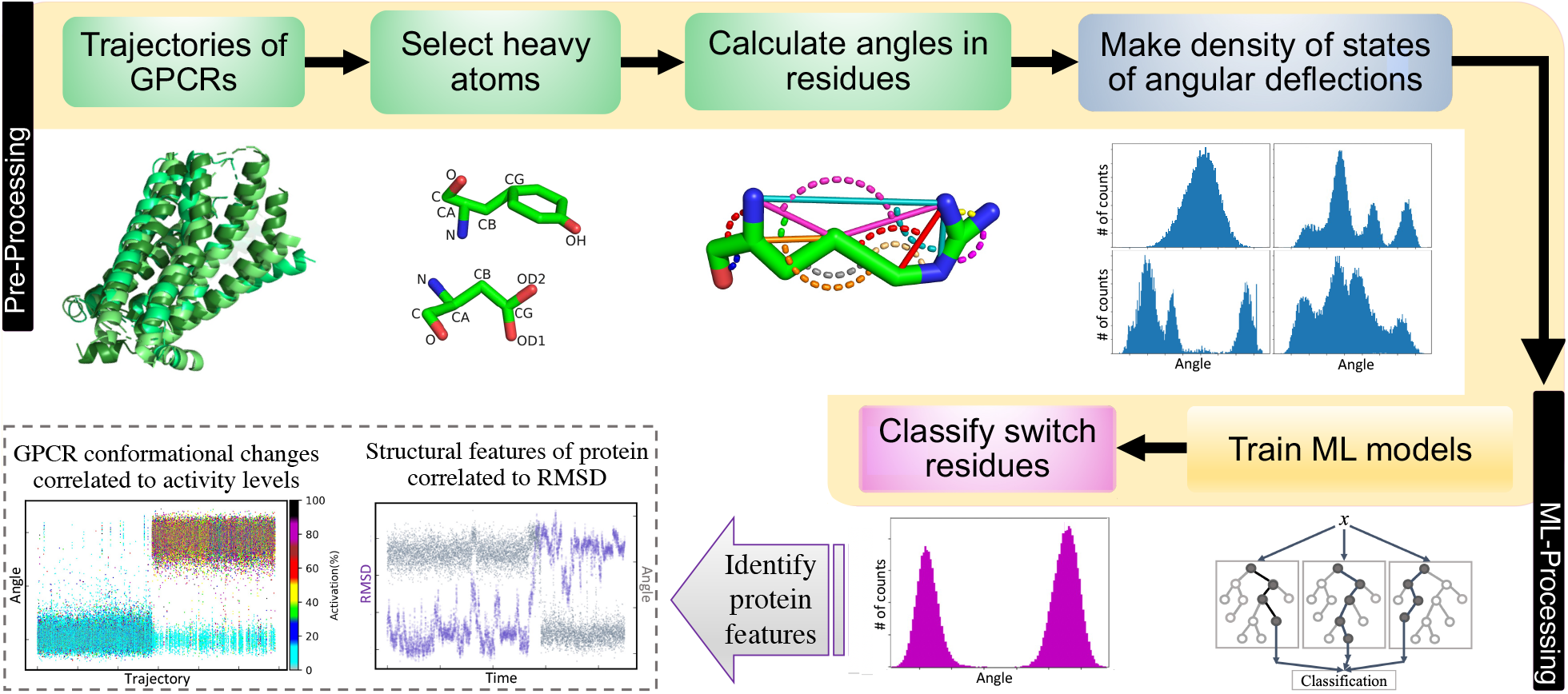
Framework of classifying bimodal-switch residues overall a protein. It is followed by two examples of using this method in Fs peptide protein and *β*_2_*AR* receptor.

**Figure 2:**
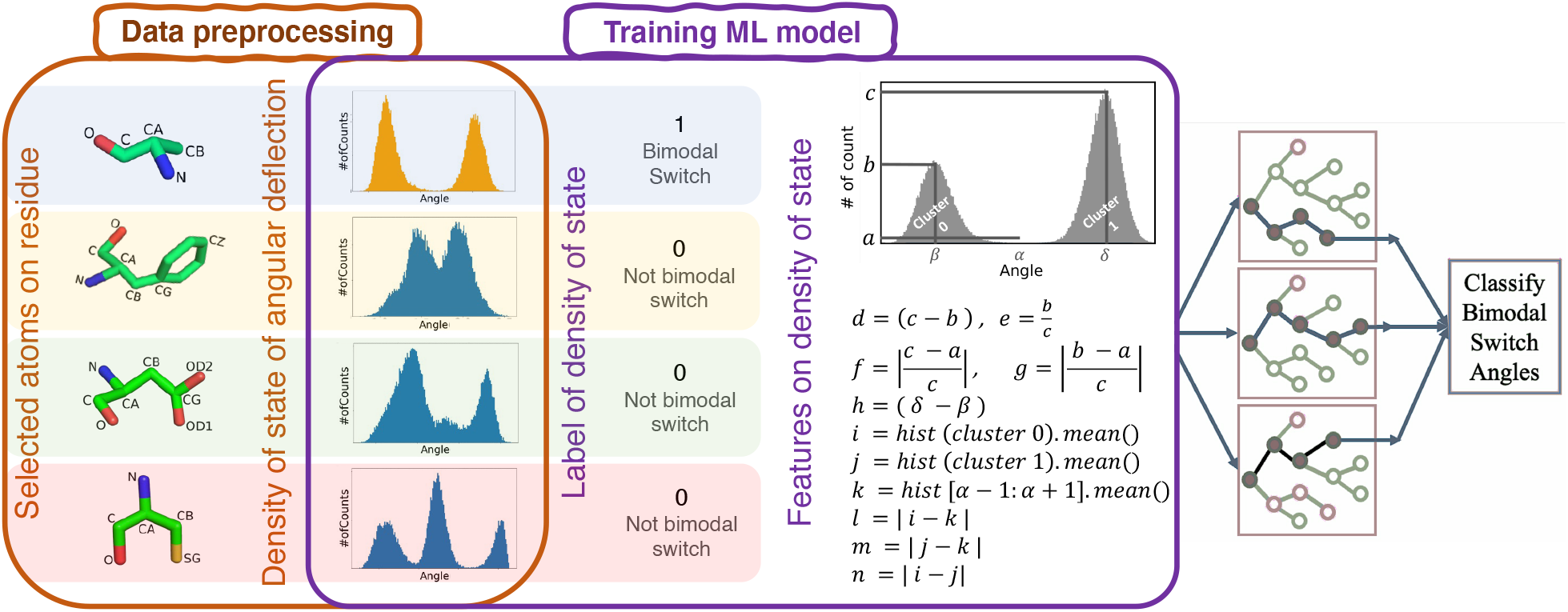
The process of data preprocessing and training shallow ML models for angular deflection classification task.

### Training ML models

In order to train an ML model capable of classifying switch angles within a protein, we used the density of states for angles as a dataset containing angle information. After obtaining various types of angle distributions, we labeled them by visual inspection that determines whether a given density represents a bimodal switch or not (see Supporting Information). As shown in Fig.2, the label of bimodal switches are 1, and others are marked as 0. Next, we defined certain features on the histograms. Since the main goal of this study is to categorize the bimodal-switch density of states which has only two distinct states without any intermediate state, we employed Kmeans algorithm with two clusters from scikit-learn library. After identifying the elements of each cluster, we proceeded to define 14 features on the density of states as defined in Fig.2. Ultimately, the ML model is trained using both the features and labels of the density of states. Through this training approach, the Random Forest model achieved a higher accuracy rate of 96.94% compared to the XGBoost and Decision Tree models (Table 1).

**Table 1:** The performances of Random Forest, XGBoost, and Decision Tree models in bimodal-switch classification tasks with standard deviations in parenthesis.

| ML Model | bimodal-switch classification accuracy (%) |
| --- | --- |
| Random Forest | 96.94 (0.04) |
| XGBoost | 96.42 (0.03) |
| Decision Tree | 95.91 (0.03) |

## Results and discussion

Proteins can undergo conformational changes occurring at different levels of protein organization. At the atomic level, conformational changes refer to the reorientation of individual atoms within a protein. The significance of atomic-level conformational changes in proteins lies in their ability to regulate protein function. We can monitor atomic-level conformational changes in amino acids by investigating angles between atoms in residues. The primary focus of this study is on bimodal-switch residues with no intermediate state existing between the two states. The bimodal-switch residues can be classified into two categories: bimodalstable-switch (BSS) and bimodal-unstable-switch (BUS) residues. We have characterized BSS residues as those that undergo a single jump in angle once a change in a global feature occurs during protein dynamics. Hence, there will be a strong correlation between the switch angle in BSS residue to the global feature in protein. In contrast, the BUS residue contributes to minor features in the protein while fluctuating between the two angles throughout the protein dynamics. To confirm our concept, we assessed the switch residues of both fs peptide protein and *β*_2_*AR* receptor by examining various global features that included BSS and BUS residues (shown in Fig.3 and Fig.4).

**Figure 3:**
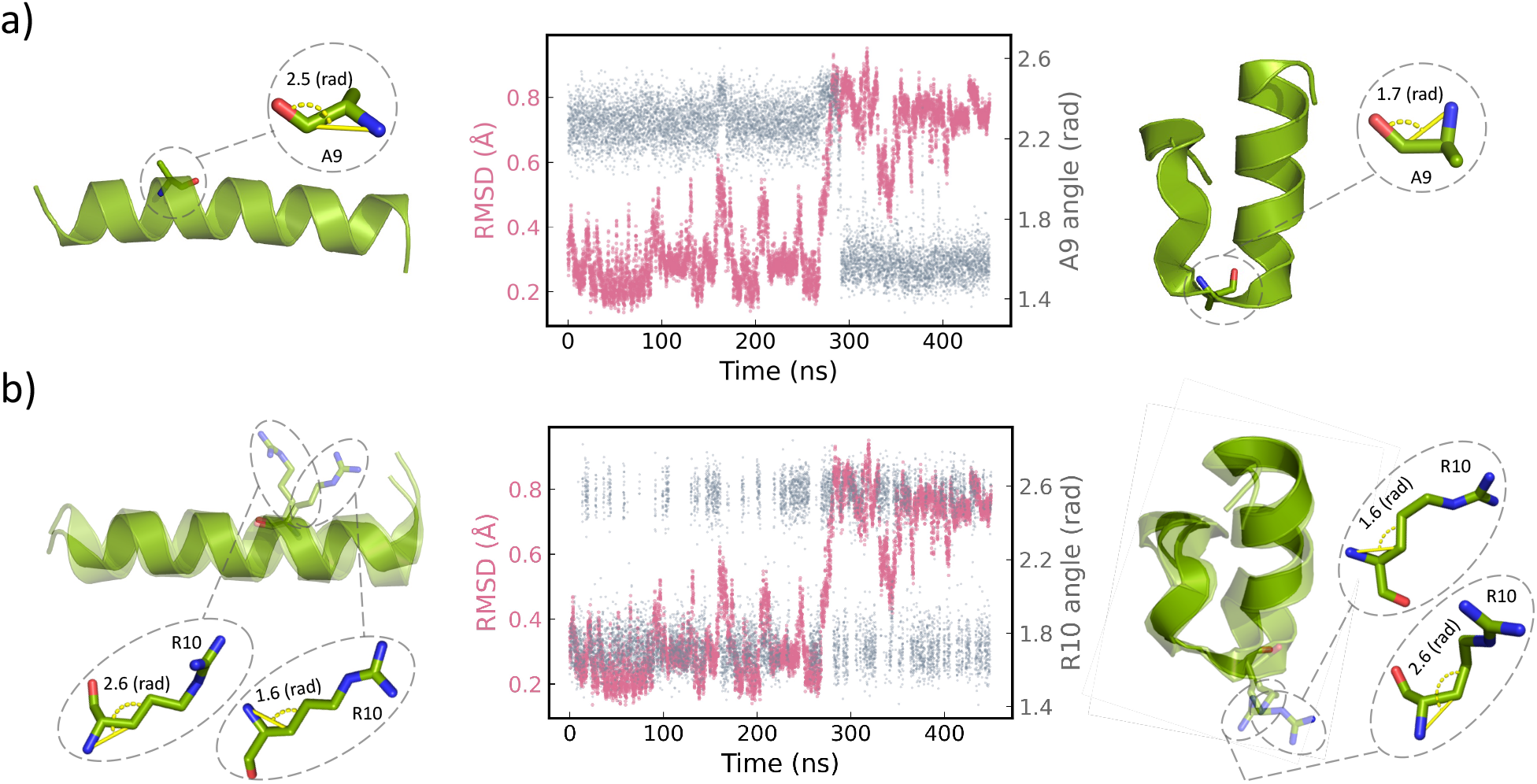
The A9 and R10 residues in Fs peptide protein classified as bimodal-switch residues using the RF model. (a) the A9 residue is a BSS residue and exhibits a strong correlation with the RMSD. (b) the R10 is a BUS residue and mostly oscillate between the two angles throughout the protein dynamics. Angular deflections in the residues are shown in PyMOL representation.

**Figure 4:**
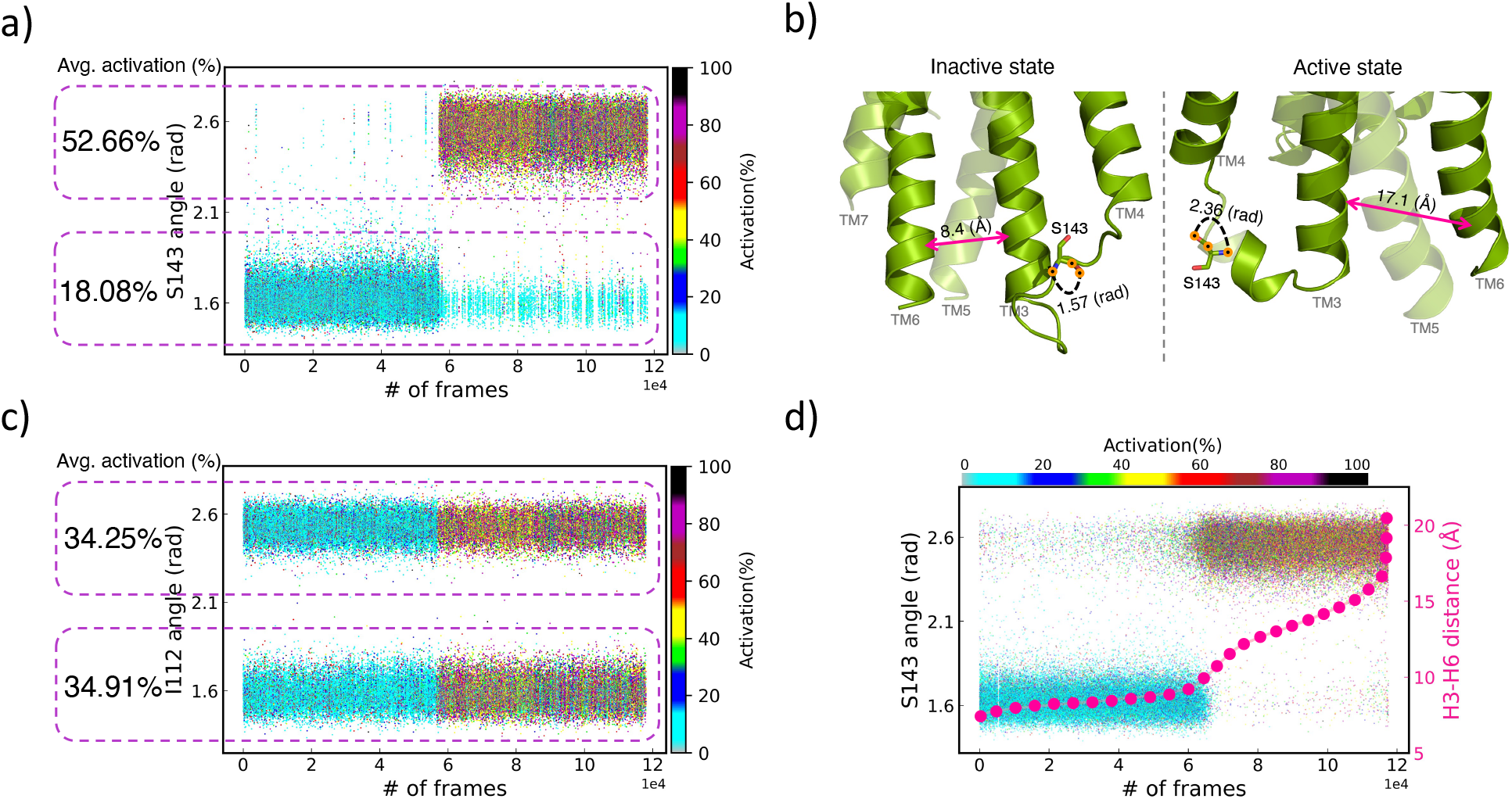
The *S*143^34.55^ and *I*112^3.31^ residues in *β*_2_*AR* receptor classified as bimodal-switch residues using the RF model. (a) the strong correlation between the *S*143^34.55^ bimodal-switch residue (as BSS residue) and activation states of the receptor. (b) the PyMOL representation of *β*_2_*AR* receptor in the active and inactive states showing the H3-H6 distance and switch angle in *S*143^34.55^ residue. (c) *I*112^3.31^ bimodal-switch residue presenting BUS function that fluctuates between the two angular states throughout the protein dynamics. (d) correlation between the *S*143^34.55^ switch residue, activation states, and H3-H6 distance in *β*_2_*AR* receptor

### Structural features of proteins correlated to RMSD

One approach to residue motion analysis is to use molecular dynamics simulations, which involve computer simulations of the protein’s behavior over time. ^36,50,51^ By analyzing the trajectories, we can identify which residues are undergoing significant conformational changes and how these changes contribute to the protein’s overall function. In this study, we used trajectories of the Fs peptide (Ace-A 5(AAARA) 3A-NME), which is a well-established model system for studying protein folding. Each trajectory is 500 nanoseconds in duration and is saved every 50 picoseconds. The simulations were executed using OpenMM 6.0.1 and the AMBER99SB-ILDN force field with GBSA-OBC implicit solvent at 300K. The simulations were initiated from randomly selected conformations obtained from an initial 400K unfolding simulation^53^ (see Supporting Information). In this trajectory, the RF model classified six residues (A3, A7, A9, R10, R15, A22) as the bimodal switch residues out of 21 residues present in the protein (shown in Table 2). The angle switch ratio (ANSR) introduces the ratio of angles identified as bimodal switch angles in a residue. For example, the A9 residue provides 10 angles in total and two of them (angles between O-C-N, and O-CA-N atoms) provide bimodal-switch angles in this trajectory, so the ANSR for A9 residue will be 2/10. The atom switch contribution (ATSC) highlights the contributions of each atom within a residue toward its switching behavior. For example, the two switch angles in A9 residue are formed between the atoms 0-C-N and O-CA-N, thus, the ATSC for atoms O, C, CA, and N will be 2,1,1, and 2, respectively. The ATSC can indicate the degree of contribution of individual atoms within a residue to its switching function. It is reasonably assumed that a residue with a high ANSR would have a strong correlation with global features in a protein. The amino acids that represent the BSS residue are highlighted in the list of switch residues presented in Table 1 and Table 2. Table 1 shows the A9 residue has the highest ANSR in the list of switch residues and is identified as a BSS residue. In this study, we specifically focused on the folding process of Fs peptide protein, as indicated by Root Mean Square Deviation (RMSD) feature.

**Table 2:** List of residues in Fs peptide protein classified as switch residues using RF model. Highlighted residue also displays a bimodal-switch-angle behavior. The ANSR presents the contribution of all angles in a residue in the switch

| Residue | ANSR | ATSC | Residue | ANSR | ATSC |
| --- | --- | --- | --- | --- | --- |
| ALA9 | 2/10 | O:2 , N:2, C:1 , CA:1 | ARG10 | 6/120 | CG:5, CB:4, C:3, N:2, NE:2, CA:1, CD:1 |
| ALA7 | 1/10 | O:1, C:1, N:1 | ARG15 | 3/120 | CG:3, C:2, CB:2, CA:1, NE:1 |
| ALA3 | 1/10 | O:1, CA:1, N:1 | ARG20 | 1/120 | CB:1, CD:1, NE:1 |
| ALA22 | 1/10 | O:1, C:1, N:1 |  |  |  |

**Table 3:**
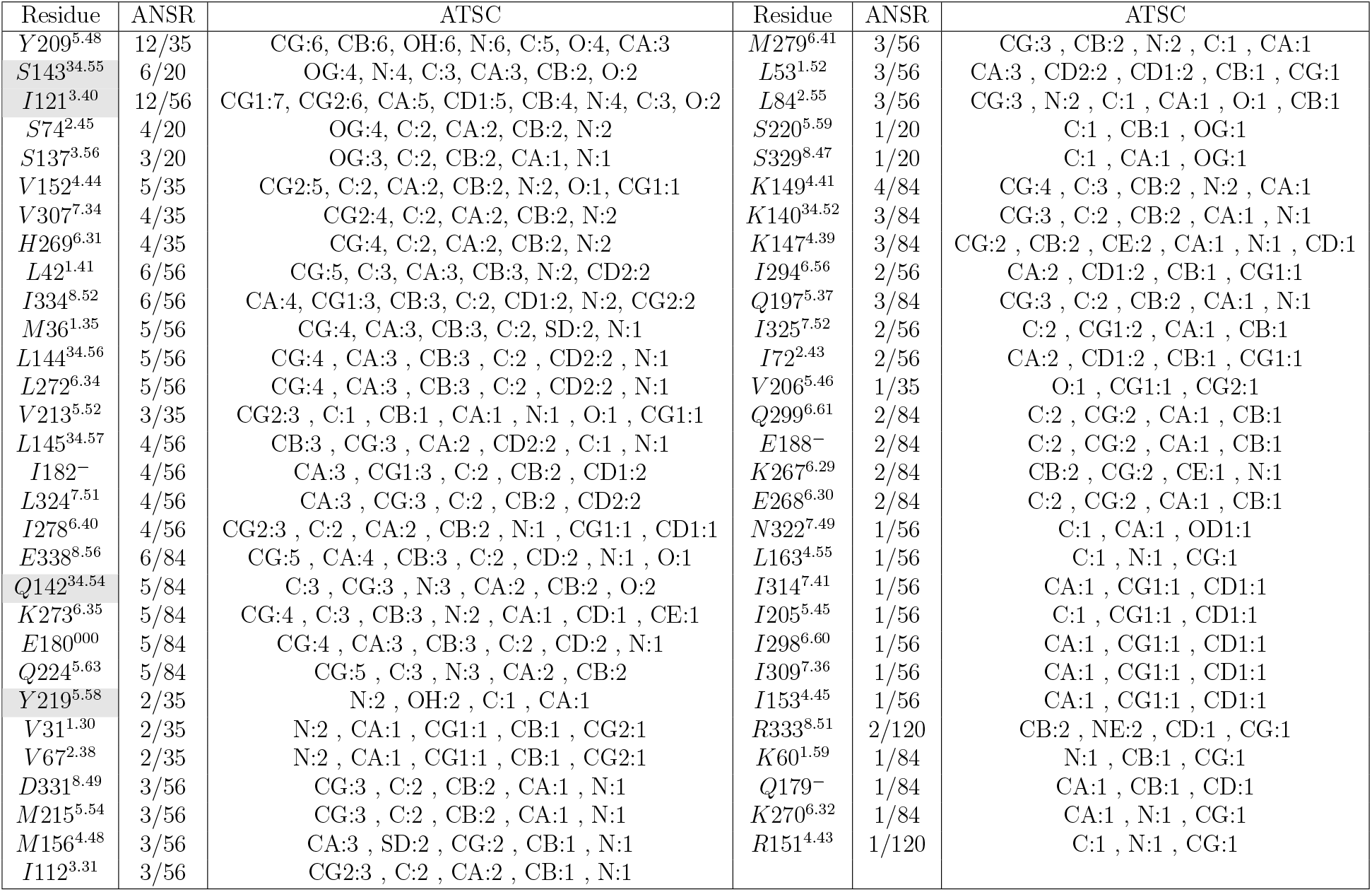
List of residues in Fs peptide protein classified as switch residues using RF model. Highlighted residue also displays a bimodal-switch-angle behavior. The ANSR presents the contribution of all angles in a residue in the switch. Ballesteros-Weinstein numbers are used for presenting the residue

After identifying the BSS residue in the Fs peptide protein, we proceeded to assess the correlation between the residue and the RMSD feature. As shown in Fig.3a, there is a strong correlation between the BSS residue and the RMSD feature in Fs peptide protein. As long as the average of RMSD is ∼ 0.3°A the average of angle between O-C-N atoms in A9 is ∼ 2.3(rad) and it switches to ∼ 1.6(rad) once the average of RMSD jumps to ∼ 2.5°A. However, the BUS residues mostly oscillate between the two angles throughout the folding process of the Fs peptide protein. Fig.3b shows the angle between O-C-N atoms in R10 residue mostly oscillate between ∼ 1.7(rad) and ∼ 2.6(rad) regardless of the significant jump in RMSD (at t ∼ 280 ns). Through such atomic-level structural analysis, we are able to investigate causalities and correlations between the small conformational changes and their contributions to the large-scale structural features in proteins. Hence, the atomic-level conformational changes are essential for regulating protein function and allowing proteins to carry out their biological roles.

### G protein-coupled receptors conformational changes correlated to activation states

G protein-coupled receptors (GPCRs), are a class of membrane proteins that are involved in many physiological processes in the body. The motions of amino acids in the ligand-binding pocket of the receptor are critical for enabling the receptor to recognize and bind to specific ligands, such as hormones, neurotransmitters, or drugs.^1,28,54–56^ Many drugs act by binding to GPCRs and modulating their activation states. After the ligand binding, GPCRs undergo a complex series of conformational changes that trigger downstream signaling pathways and enable interaction with a G protein. The motions of amino acids in the transmembrane and intracellular regions of the receptor are critical for transmitting these conformational changes and initiating downstream signaling events. There are two main activation states of GPCRs: inactive and active. In the inactive state, the receptor is not bound to a ligand and is not interacting with a G protein. In this state, the receptor has a low affinity for G proteins. The ability of GPCRs to switch between different activation states allows for the precise control of downstream signaling pathways and ultimately determines the physiological response of the cell. One important significance of the activation states of GPCRs is their involvement in signal transduction pathways allowing cells to respond appropriately to drugs targeting the receptor. GPCRs interact with G proteins to initiate downstream signaling events.

The motions of amino acids in the intracellular regions of the receptor are important for facilitating this interaction and inducing conformational changes in the G protein.^1,28,55–58^ Understanding conformational changes in small and large scales of GPCRs is important for developing new therapeutics, ligand binding, signal transduction, and G protein activation. The switch residues investigation in GPCRs can provide insight into the conformational changes and the transition pathways between activation states in the receptors. In this study, we analyzed switch residues in *β*_2_*AR* receptor that plays a crucial role in regulating bronchodilation, heart rate, and blood pressure.^59–61^ We selected 5000 trajectories randomly from the numerous short trajectories of the receptor in the presence of partial inverse agonist carazolol (PDB 3P0G),^52,62^ to assess the switch angles and correlation between BSS residue to the receptor’s activation states.

Table 2 shows list of residues in this receptor that are classified as switch residues using the RF model. Out of 12138 angles generated by 271 amino acids in *β*_2_*AR* receptor, the RF model classifies 171 switch residues (see Supporting Information). Those switch residues are created by 59 unique amino acids, as shown in Table 2. Similar to Table 1, the highlighted residues identified as BSS residues. In our previous work,^57^ we introduced XGBoost model that could predict the activation state with prediction accuracy of 97.27% and 8.55% MAE for activation regression. We applied the model on the dataset employed to classify switch angles, with the purpose of assessing the correlation between the activation states and the switching function of residues. Fig4a demonstrates that there is a strong correlation between the activity states and the *S*143^34.55^ switch residue. As the *S*143^34.55^ angles range from ∼ 2.25 *−* 2.75 (rad), the average activity level of the receptor is 52.66%. Conversely, when the *S*143^34.55^ switch angle range from approximately ∼ 1.45*−*1.80 (rad), the average activity level of the receptor drops significantly to 16.47%. However, some of the switch residues have no significant relation to the activation states of the receptor. As shwon in Fig.4c, the *I*112^3.31^ switch angle mostly fluctuate between the two angular states and the average of activation states of the protein changes between 34.25% and 34.91% during the protein dynamics. To verify the performance of the ML model predicting the activity level of a given receptor, we incorporated the helix3-helix6 (H3-H6) distances to this result. Helix3 and helix6 are two of the seven transmembrane helices that make up the core structure of GPCRs. Changes in this distance can modulate the conformation of the receptor and affect downstream signaling pathways, receptor activation, desensitization, and internalization making it a critical determinant of receptor function. The H3-H6 distance is measured as *C*_*α*_ contact distance between *R*131^3.50^-*L*272^6.34^ amino acids. Fig.4b represents a snapshot of the *β*_2_*AR* receptor in the active and inactive states presenting the H3-H6 distance and the switch angle in *S*143^34.55^ residue. As shown in Fig.4d, the H3-H6 distance remarkably changes in the transition between the two angles which is heavily correlated to the activation states of the receptor.

## Conclusion

Even small changes in the atomic-level conformation of a residue can have a significant impact on protein activity and function. In this study, we examined the reorientation of individual atoms within a protein by analyzing the angles between the atoms present in each residue throughout the entire protein. We mainly focused on bimodal switch resides that can transition between two distinct angular states with no intermediate states in between. We trained shallow ML models to classify the bimodal-switch residues with features on the density of states of angular features. Our analysis indicates that the Random Forest model was particularly effective, achieving a remarkable accuracy of 96.94% in identifying bimodal-switch residues. Our study has revealed that bimodal-switch residues can be classified into two categories: bimodal-stable-switch (BSS) and bimodal-unstable-switch (BUS) residues. The BSS residues tend to remain stable throughout the protein dynamics and only switch between the two angular states once a change in a global feature of the protein occurs. On the other hand, the BUS residues primarily fluctuate between the two angular states during the protein dynamics, regardless of significant changes in the global protein features. We validated our concepts by analyzing the Fs peptide protein and *β*_2_*AR* receptor. Our results revealed a strong correlation between a BSS residue (A9) and the protein folding process, as indicated by RMSD values observed in the Fs peptide protein. Additionally, we found a significant correlation between BSS residues in the *β*_2_*AR* receptor (*S*143^34.55^, *I*121^3.40^, *Q*142^34.54^, *Y* 219^5.58^) and its activation states. Understanding these types of conformational changes is vital for analyzing protein structure, activity, stability, and function. Moreover, it is a crucial aspect in the design of drugs targeting specific protein conformations, especially in the case of GPCRs, where even small conformational changes can have significant impacts on both the receptor’s downstream signaling pathways and the drug’s efficacy.

## Acknowledgement

The authors gratefully acknowledge the use of MD simulation trajectories of *β*_2_*AR* receptor dataset provided by Stanford University.^62^ This work is supported by the Center for Machine Learning in Health (CMLH) at Carnegie Mellon University and a start-up fund from Mechanical Engineering Department at CMU.

## Notes

### Competing Interest Statement

The authors have declared no competing interest.

